# FTY720 (Fingolimod), a modulator of sphingosine-1-phosphate receptors, increases baseline hypothalamic-pituitary adrenal axis activity and alters behaviors relevant to affect and anxiety

**DOI:** 10.1101/2021.05.10.443108

**Authors:** Brian Corbett, Sandra Luz, Nathaniel Sotuyo, Jiah Pearson-Leary, Ganesh S. Moorthy, Athena F. Zuppa, Seema Bhatnagar

**Affiliations:** Center for Stress Neurobiology, Children’s Hospital of Philadelphia, Philadelphia, Pennsylvania, USA; Perelman School of Medicine, University of Pennsylvania, Philadelphia, Pennsylvania, USA; Center for Clinical Pharmacology, The Children’s Hospital of Philadelphia, Philadelphia, Pennsylvania, USA; Department of Anesthesiology and Critical Care Medicine, The Children’s Hospital of Philadelphia, Philadelphia, Pennsylvania, USA

**Keywords:** stress, sphingosine-1-phosphate, S1PR, HPA axis, angiogenesis, FTY720, medial prefrontal cortex

## Abstract

FTY720 (fingolimod) is an analog of sphingosine, a ubiquitous sphingolipid. Phosphorylated FTY720 (FTY720-P) non-selectively binds to sphingosine-1-phosphate receptors (S1PRs) and regulates multiple cellular processes including cell proliferation, inflammation, and angiogenesis. We recently demonstrated that S1PR3 expression in the medial prefrontal cortex (mPFC) of rats promotes stress resilience and that S1PR3 expression in blood may serve as a biomarker for PTSD. Here we investigate the effects of FTY720 in regulating the stress response. We found that single and repeated intraperitoneal injections of FTY720 increased baseline plasma adrenocorticotropic hormone (ACTH) and corticosterone concentrations. FTY720 also mitigated restraint-induced increases in ACTH and corticosterone. FTY720 reduced social anxiety- and despair-like behavior as assessed by increased social interaction time and reduced time spent immobile in the Porsolt forced swim test. In blood, FTY720 administration reduced lymphocyte and reticulocyte counts, but raised erythrocyte counts. FTY720 also reduced mRNA of *angiopoietin 1*, *endothelin 1*, *plasminogen 1*, *Vegf-B*, and *Mmp2* in the medial prefrontal cortex, suggesting that FTY720 reduced angiogenesis. The antidepressant-like and anxiolytic-like effects of FTY720 may be attributed to reduced angiogenesis as increased stress-induced blood vessel density in the brain contributes to depression- and anxiety-like behavior in rats. Together, these results suggest that S1PRs regulate baseline HPA axis activity but reduces social anxiety and despair providing further evidence that S1PRs are important and novel regulators of stress-related functions.

## 1. INTRODUCTION

Sphingosine-1-phosphate receptors (S1PRs, also known as endothelial differentiation genes or *edg* receptors) are a class of G-protein coupled receptors (GPCRs) that bind the endogenous ligand spingosine-1-phosphate (S1P). The S1PRs regulate a number of cellular processes including inflammation, angiogenesis, migration, differentiation, and proliferation in peripheral organs and blood cells ^1–7^. Little is known about their functions in the brain or on brain-regulated processes^8–10^. We recently reported that S1PR3 expression in the medial prefrontal cortex (mPFC) promotes resilience to the adverse effects of stress by mitigating stress-induced TNFα production in the mPFC ^8^. Furthermore, S1PR1 activation in the hippocampus can increase adult neurogenesis ^9^. These studies highlight the importance of S1PRs in the brain for mediating multiple functions relevant to our understanding of affect and anxiety disorders.

FTY720 (fingolimod) is an FDA-approved pharmaceutical (Gilenya) that is currently used for the treatment of relapsing-remitting multiple sclerosis (MS) and was the first orally administered disease-modifying therapy for long term MS treatment ^10,11^. As a structural analog of S1P, FTY720 modulates S1PR activity following its phosphorylation by sphingosine kinase 2 into FTYP ^12,13^. FTY720 targets S1PRs non-specifically, binding to S1PRs1,3-5 with a particularly high affinity for S1PR1 ^14–16^. Although FTY720 is an agonist at S1PR1 initially, chronic administration of FTY720 causes internalization of S1PR1s resulting in functional antagonism of S1PR1 ^16,17^. Functional antagonism of S1PR1 on lymphocytes is thought to underlie the efficacy of FTY720 in treating MS as it prevents lymphocytes from leaving lymph nodes, thereby reducing the autoimmune response ^17,18^. However, FTY720 does not cause internalization of S1PR3 and serves as a partial agonist of S1PR3 ^16^. Therefore, FTY720 does not modulate the activity of all S1PRs through a common mechanism

Beyond these immunosuppressive effects of FTY720, little is known about its ability to regulate behavior and physiology. FTY720 reduces immobility in the Porsolt forced swim test (FST) suggesting reductions in behavioral despair and reduces social anxiety-like behavior as assessed by increased interaction time with a novel conspecific ^19^. FTY720-P inhibits hippocampal histone deacetylases (HDACs) ^20^ and increases hippocampal brain-derived neurotrophic factor (BDNF) ^20,21^, which may contribute to reductions in behavioral despair ^22^. Further, hippocampal neurogenesis is increased with chronic FTY720 treatment ^23^. FTY720 may also reduce inflammatory processes in the brain through partial agonism of S1PR3 ^16^. Therefore, there are multiple mechanisms by which FTY720 could modulate brain function that reduce affective- or anxiety-like behavior.

S1PRs are expressed in the hypothalamus ^24^, pituitary ^25^, and adrenal glands ^26^, and therefore may regulate hypothalamic-pituitary-adrenal (HPA) axis function. However, the role of S1PRs in modulating HPA activity is not clear. Here, we studied the effects of FTY720 on HPA axis activation and affective- and anxiety-like behaviors. Additionally, we studied cellular and molecular alterations that may contribute to these physiological and behavioral effects of FTY720. Because FTY720 reduces lymphocyte counts ^27–30^, we investigated the effects of FTY720 on counts of specific blood cell types. Additionally, functional antagonism of S1PR1 by FTY720 inhibits angiogenesis ^31^. Blood vessel plasticity is increased in rats that are vulnerable to the adverse effects of stress and contributes to their increases in behavioral despair and anxiety-like behaviors ^32^. Therefore, we investigated the expression of angiogenesis markers in the mPFC, a brain region that plays a key role in regulating stress resilience/vulnerability ^8,33^.

## 2. METHODS

### 2.1. Animals

Adult male Sprague–Dawley rats (225–250 g) were obtained from Charles River Laboratories (Wilmington, MA, USA). Rats were singly housed in polycarbonate cages with standard bedding and with food and water available *ad libitum*. Animals were acclimated to a 12-h light–dark cycle with lights on at 06:15 and lights off at 18:15 in a temperature-controlled vivarium for at least 5 days prior to administration of any stress and/or drug administration protocols. All experiments took place during the inactive phase between 1000 and 1400 h. Rats were euthanized by rapid decapitation and their brains were immediately snap-frozen in 2-methylbutane. Experiments were performed in compliance with all relevant ethical regulations for animal testing and research. Experiment protocols followed the NIH Guide for the Care and Use of Laboratory Animals and were approved by the Children’s Hospital of Philadelphia Research Institute’s Animal Care and Use Committee.

### 2.2. Experimental Procedures

#### 2.2.1. Timecourse of FTY720 and FTY720-P concentrations in plasma following ip injection

Rats were administered a single FTY720 ip injection at a dose of 2.5 mg/kg. This dose is sufficient to affect S1PR signaling ^34^ and is below higher functional doses (e.g. 10 mg/kg) that have been reported in the absence of adverse side effects ^30,35^ . FTY720 and FTY720-P concentrations in rat plasma was determined by liquid chromatography-tandem mass spectrometry analysis. FTY720 and FTY720-P were extracted from rat plasma with methanol. Electrospray ionization in the positive ion mode was utilized for the selective detection of FTY720 (m/z 308.3→ 255.2) and FTY720-P (m/z 388.2→ 290.1) using C17-sphingosine (m/z 286.2 → 238.1) as an internal standard with AB Sciex 4000 mass spectrometer. The separation was accomplished utilizing Xbridge C18 (50 × 2.0 mm id., 3.5 μm) column with Shimadzu LC 20AD HPLC system and a run time of 6.0 min. The assay was linear over the range of 0.5 to 250 ng/mL for FTY720 and 1.0 to 500 ng/mL FTY720-P in rat plasma. Rat plasma samples (50 μL) at different time points were analyzed to determine the concentration of FTY720 and FTY720-P. Group sizes were n = 3/group, standard in mass spectroscopic analyses.

#### 2.2.2. Effects of acute peripheral administration of FTY720

##### 2.2.2.1. Effects of acute peripheral administration of FTY720 on basal and acute restraint-induced HPA axis activity

Rats were administered a single injection of saline or FTY720 (2.5 mg/kg, i.p.). Sixty min following injection, rats were restrained for 30 min in a plexiglass tube (20.7 cm long × 11 cm in diameter). Tail blood was taken (~400 μL) at 0min (baseline), and at 15min and 30min (during restraint). Rats were returned to their home cage and 30min later (at the 60min timepoint) trunk blood samples were collected (recovery timepoint).Radioimmunoassay kits from MP Biomedical (Orangeburg, NY, USA) were used to assay plasma samples for adrenocorticotropic hormone (ACTH) and corticosterone. Two independent cohorts of rats were used. Group sizes were n=10/group.

##### 2.2.2.2. Effects of repeated peripheral FTY720 administration on basal HPA axis activity

We examined the effects of steady-state concentrations of FTY720 achieved by 3 daily injections of saline or FTY720 (2.5 mg/kg, ip) administered 24 hours apart on plasma ACTH (saline n=5, FTY720 n=8) and corticosterone (saline n=5, FTY720 n=10) concentrations. Blood samples were collected 60min after the third injection.

### 2.3. Effects of repeated peripheral administration of FTY720 on specific blood cell type populations

Rats received daily injections of saline or FTY720 (2.5 mg/kg, i.p.) for three days. Sixty minutes following the third injection, 50 μL of blood was collected and added to a collection tube containing 10 μL of ethylenediaminetetraacetic acid (EDTA). Blood samples were diluted 1:5 with Sysmex CELLPACK (EPK, sodium chloride 6.38 g/L, boric acid 1.0 g/L, sodium tetraborate 0.2 g/L, EDTA-2k 0.2 g/L) and underwent hematological analysis conducted by the Children’s Hospital of Philadelphia Hematology Core using the Sysmex XT-2000i. The Sysmex XT-2000i uses the electric resistance detecting method with hydrodynamic focusing to measure red blood cell shape and volume, hematocrit, mean platelet volume, and mean corpuscular volume. A 633 nm semi-conductor laser is used for flow cytometry analysis of the quantities of specific blood cell types including erythrocytes, reticulocytes, lymphocytes, monocytes, total white blood cells, neutrophils, eosinophils. Side scatter is sued to determine the internal complexity of the cell (size, shape, nuclear/granule densities). Hemoglobin is measured photocolorimetrically using SLS-HGB. Group sizes were: saline n=7, FTY720 n=8.

### 2.4. Effects of repeated administration of FTY720 on behaviors

#### 2.4.1. Effects on repeated peripheral FTY720 treatment on behavior in the social interaction test

Rats received daily injections of saline or FTY720 (2.5 mg/kg, i.p.) for three days. Sixty minutes following the third injection, rats were subjected to the social interaction test. Rats were placed in an open field black box (90 cm × 90 cm) with an age-matched novel stimulus rat of the same strain (Sprague-Dawley) and of a similar size and allowed to interact for 15min. Time interacting with the stimulus rat was defined by the time the experimental rat was actively investigating the stimulus rat with its snout closer than 3 cm away (approximately the length of the snout of the rat) from the stimulus rat. Each interaction was videotaped and coded for social interaction time by 2 coders who were blind to the experimental conditions. Group sizes were: saline n=5, FTY720 n=6.

#### 2.4.2. Effects of repeated peripheral FTY720 treatment on behaviors in the open field

Rats received daily injections of saline or FTY720 (2.5 mg/kg, i.p.) for three days. Sixty minutes following the third injection, rats were subjected to the open field test. Rats were placed alone in an open field black box (90 cm × 90 cm) and video recorded for 15 minutes. The Noldus EthoVision^®^XT software automated tracking and analysis function was used to analyze the total distance traveled during the 15min trial. Group sizes were n=9/group.

#### 2.4.3. Effects of repeated peripheral FTY720 treatment on behaviors in the Forced Swim Test

Rats were placed in a glass cylinder filled with 60 cm of water so that their tails could not touch the bottom of the cylinder while floating. Rats underwent a 15 min training phase followed by a 5 min test phase on the following day. Similar to previous studies assessing the efficacy of antidepressants ^36^, rats were treated with FTY720 (2.5 mg/kg, ip) immediately following the training phase and 60 minutes prior to the test phase. The test phase was videotaped and coded for time engaged in immobile, swimming, and climbing behaviors by 2 coders who were blind to the experimental conditions. Group sizes were n=8/group.

#### 2.4.4. Effects of centrally administered FTY720 on behaviors in the Porsolt FST and brain concentrations

We next examined whether the behavioral changes induced by peripheral administration of FTY720 were due to direct effects of FTY720 in the brain. Rats were anesthetized with a cocktail of ketamine, xylazine, and acepromazine (5:0.1:1, 1 μL/g) and fitted with intracerebral cannulae to target the lateral cerebral ventricles (A/P: Bregma −1.1 mm, M/L: ± 1.5 mm, D/V −3.2 mm). Three additional holes were drilled and bone screws (Plastics One, 0-80) were fastened in the skull (Bregma – 3.5 mm, 5.0 mm left; Bregma – 2.0 mm, 4.0 mm right; Bregma – 8.0 mm, 6.0 mm right) to securely fix the cannulae in place. The bilateral cannulae (Plastic One, C232I/SPC), which allow for a 1 mm projection ventral to their placement, were positioned 2.2 mm ventral to the skull surface. Dental cement was used to adhere the cannulae to the skull and bone screws. 22 gauge double dummy cannulae (Plastics One, C232DC/spc) were placed in the intracerebral cannulae to prevent the cannulae from closing. An infusion dust cap (Plastics One, 303DC/1) was used to cover the cannulae and fasten the dummy cannulae. Rats were administered meloxicam (300 μL) and monitored until they woke up and were allowed to recover for 7 days. For each experiment, the dummy cannulae were removed and a hydraulic syringe filled with saline was used to inject 1 μL of either saline or FTY720 (1 μg/μL) in awake rats. Dummy cannulae were replaced and the rats were returned to their homecage or underwent behavioral testing. Similar to methods assessing the efficacy of antidepressant drugs ^53^, we administered FTY720 immediately after the 15-minute training phase of the FST and 60 minutes prior to the 5-minute FST test phase. Punches (1mm) of the basolateral amygdala (BLA), medial prefrontal cortex (mPFC) and dorsal and ventral hippocampal (dHPC and vHPC) regions were collected at 60 min following icv administration of FTY720. Concentrations of FTY720 and FTY720-P were assessed in both plasma and these brain regions by liquid chromatography-tandem mass spectroscopy as in the first experiment. Group sizes were: saline n=6, FTY720 n=10.

### 2.5. Effects of repeated peripheral administration of FTY720 on expression of angiogenesis markers in the mPFC

Rats received daily injections of saline or FTY720 (2.5 mg/kg, i.p.) for three days. Sixty minutes following the third injection, rats were sacrificed and their brains were snap-frozen in 2-methylbutane. Snap-frozen brains were placed on a cryostat and the mPFC was dissected using a standard punch method. RNA was isolated using the RNeasy Plus MicroKit (74034, Qiagen) and carried out per the manufacturer’s instructions. Approximately 500 ng of RNA was collected per mPFC sample. RNA was used as a template for cDNA synthesis using the RT^2^ First Strand Kit (330404, Qiagen). A customized RT2 Profiler PCR array for rat angiogenesis markers (PARN 042Z, 330231, Qiagen) containing SYBR Green RT PCR assays for 84 genes of interest, and 3 synthetic control genes (reverse transcription control, positive PCR control and rat genomic DNA contamination control) was used to run quantitative PCR on an ABI 7500 PCR system. β2-microglobulin was used as a housekeeping gene. Samples were run according to manufacturer’s instructions. The comparative Ct method was used to plot mRNA expression differences for genes of interest. Ct values were normalized to the average Ct values of the five housekeeping genes for each rat. Group sizes were: saline n=7, FTY720 n=8.

### 2.6. Statistical Analyses

Statistical analyses were performed using Prism 5 and 7. Differences between means of two groups were assessed using a two-tailed, unpaired Student’s t test unless otherwise indicated. One-way ANOVA was used to analyze FTY720 and FTY720-P plasma concentrations following a single FTY720 injection. Repeated-measures two-way ANOVA (Drug × Time, with repeated measures on Time) with Tukey’s multiple comparisons post hoc tests were used for analyzing data over multiple time points (e.g. restraint-induced ACTH and corticosterone concentrations) when comparing saline- and FTY720-treated rats. *p* values were set at p<0.05. Data beyond three standard deviations from the mean were considered outliers and discarded from analysis.

## 3. RESULTS

### 3.1. Timecourse of FTY720 and FTY720-P concentrations in plasma following intraperitoneal (ip) administration

We observed significant changes in concentrations of FTY720 (F_11,24_ = 9.72, p < 0.0001) and FTY720-P (F_11,24_ = 19.6, p < 0.0001) over time in plasma following a single injection of FTY720 (2.5 mg/kg, ip). Compared to baseline (0min) values, FTY720 was increased at 15 min (p = 0.009), 30 min (p = 0.04), 1 hour (p = 0.002), 4 hours (p = 0.0031), 8 hours (p = 0.004), and 24 hours (p = 0.0001). Compared to baseline (0min) values, FTY720-P was increased at 4 hours (p < 0.0001), 8 hours (p < 0.0001), 24 hours (p < 0.0001), 48 hours (p = 0.0002), and 72 hours (p = 0.03). These results show that FTY720 concentrations in blood are increased 30min after injection and remain elevated up to 24h later. Because of the delayed and prolonged increase in FTY720-P following a single injection, FTY720 was injected every 24 hours in experiments in which FTY720 was injected repeatedly to ensure elevated steady-state plasma concentrations of FTY720-P (Fig. 1A).

**Figure 1.**
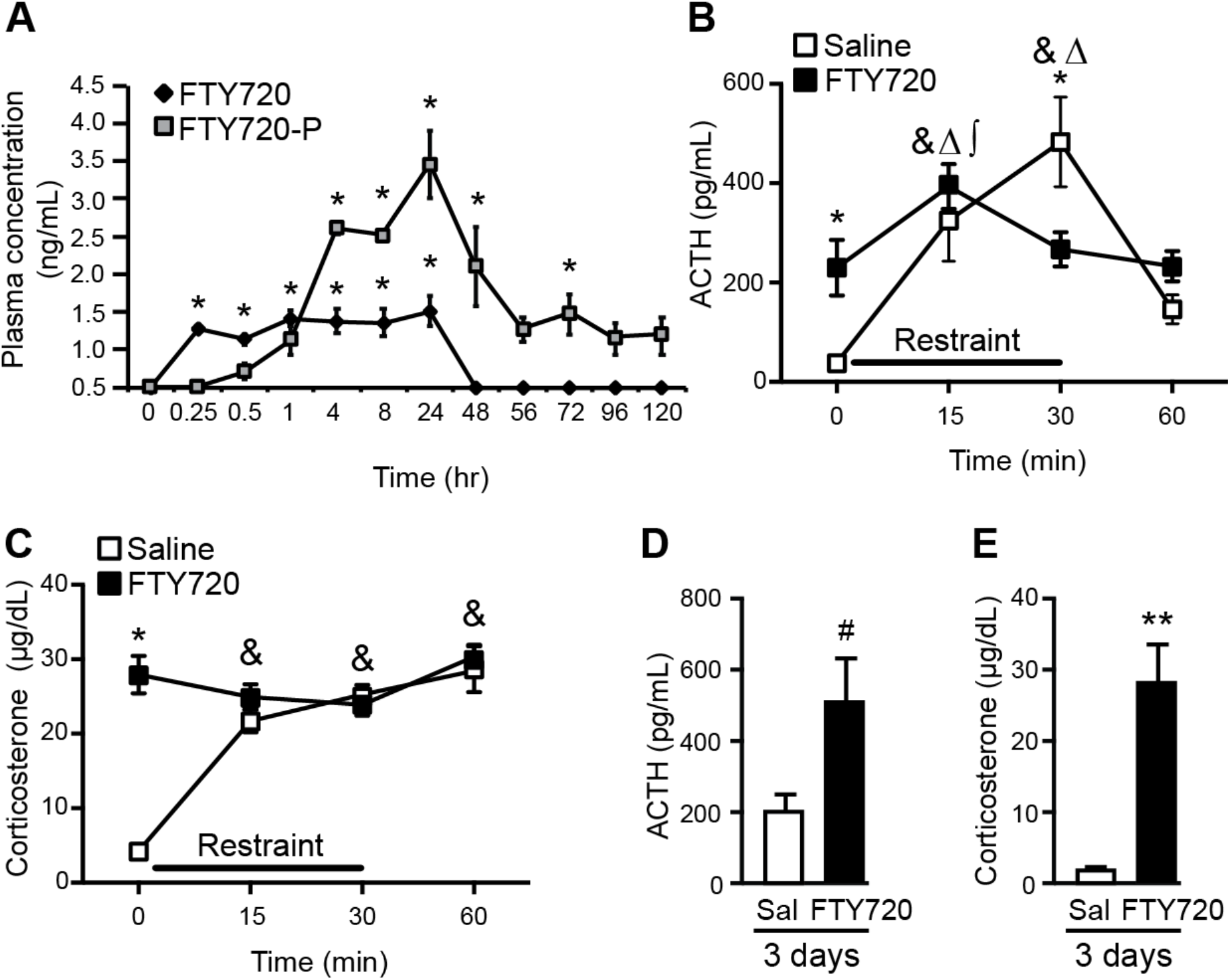
Intraperitoneal administration of FTY720 increased baseline ACTH and corticosterone concentrations. A) Characterization of plasma concentrations of FTY720 and FTY720-P by liquid chromatography-tandem mass spectrometry analysis over days following a single injection of FTY720 (2.5 mg/kg i.p.). B) Plasma ACTH concentrations in saline- and FTY720-treated rats following a single injection of FTY720. C) Plasma corticosterone concentrations in saline- and FTY720-treated rats following a single injection of FTY720. D) Basal ACTH at 60min following a third daily injection of saline or FTY720. E) Basal corticosterone at 60min following a third daily injection of saline or FTY720. Bars/squares represent group means ± SEM. For FTY720 and FTY720-P concentrations in A, * indicates significant increase compared to its respective baseline (0min) value. For B-E, **p<0.01, *p<0.05, #p<0.1 between drug groups. & denotes significant increase compared to 0min baseline in saline-treated rats. Δ denotes significant increase compared to 60min timepoint in saline-treated rats. ∫ denotes significant increase compared to 0min baseline in FTY720-treated rats.

### 3.2 Effects of acute peripheral administration of FTY720

#### 3.2.1. Effects of acute peripheral administration of FTY720 on basal and acute restraint-induced HPA axis activity

For ACTH, results of the two-way repeated measures analyses (Drug × Time) showed a significant Time effect (F_3,54_ = 15.13, p < 0.0001), and a significant Interaction effect (F_3,54_ = 7.862, p = 0.0002) but not a significant Drug group effect. Post-hoc analyses of the significant Interaction effect indicated that plasma ACTH in saline-treated rats was increased at 15min and 30min compared to 0min baseline and ACTH concentrations at 60min were reduced compared to 15min and 30min and not significantly different from 0min baseline concentration. In FTY720-treated rats, post-hoc analyses indicated a significant increase in plasma ACTH at 15min compared to 0min baseline but ACTH concentrations at 30min and at 60min were not significantly different from baseline. Comparing saline to FTY720 treatment, ACTH concentrations were higher in FTY720-treated rats at baseline (0min) compared to saline treated rats but were lower at 30min (Fig. 1B).

For corticosterone, results of the two-way repeated (Drug × Time) measures analyses showed a significant Drug Group effect (F_1,18_ = 18.7, p = 0.0004), a significant Time effect (F_3,54_ = 19.62, p < 0.0001) and a significant Interaction effect (Interaction F_3,54_ = 21.62, p < 0.0001). The significant Drug Group indicated that FTY720-injected animals had significantly increased overall concentrations of corticosterone compared to saline-injected animals. Post-hoc analyses of the significant Interaction effect indicated that plasma corticosterone concentrations in saline-treated rats were increased at 15min, 30min and 60min compared to 0min baseline. In FTY720-treated rats, no significant differences across time were observed in corticosterone concentrations. Furthermore, corticosterone concentrations were higher at 0min in FTY720-treated compared to saline-treated rats (p<0.0001) (Fig. 1C).

#### 3.2.2. Effects of repeated peripheral FTY720 administration on basal HPA axis activity

Basal ACTH concentrations trended toward an increase (t_11_ = 1.862, p = 0.089) and basal corticosterone concentrations were significantly increased (t_13_ = 3.833, p = 0.0021) in FTY720-treated rats compared to saline-treated rats (Fig. 1D,E),

### 3.3. Effects of repeated peripheral administration of FTY720 on specific blood cell type populations

Compared to vehicle-treated rats, FTY720-treated rats displayed reductions in lymphocyte counts (t_13_ = 8.59, p < 0.0001) (Fig. 2A) and in the percentage of total reticulocytes that were immature (t_13_ = 2.296, p = 0.0389) (Fig. 2B). FTY720 treatment produced an increase in the percentage of total blood volume composed of erythrocytes (t_13_ = 3.01, p = 0.01) (Fig. 2C). FTY720 treatment had no effects on the overall counts of reticulocytes, monocytes, eosinophils, platelets, neutrophils or mean corpuscular volume (Fig. 2D-I).

**Figure 2.**
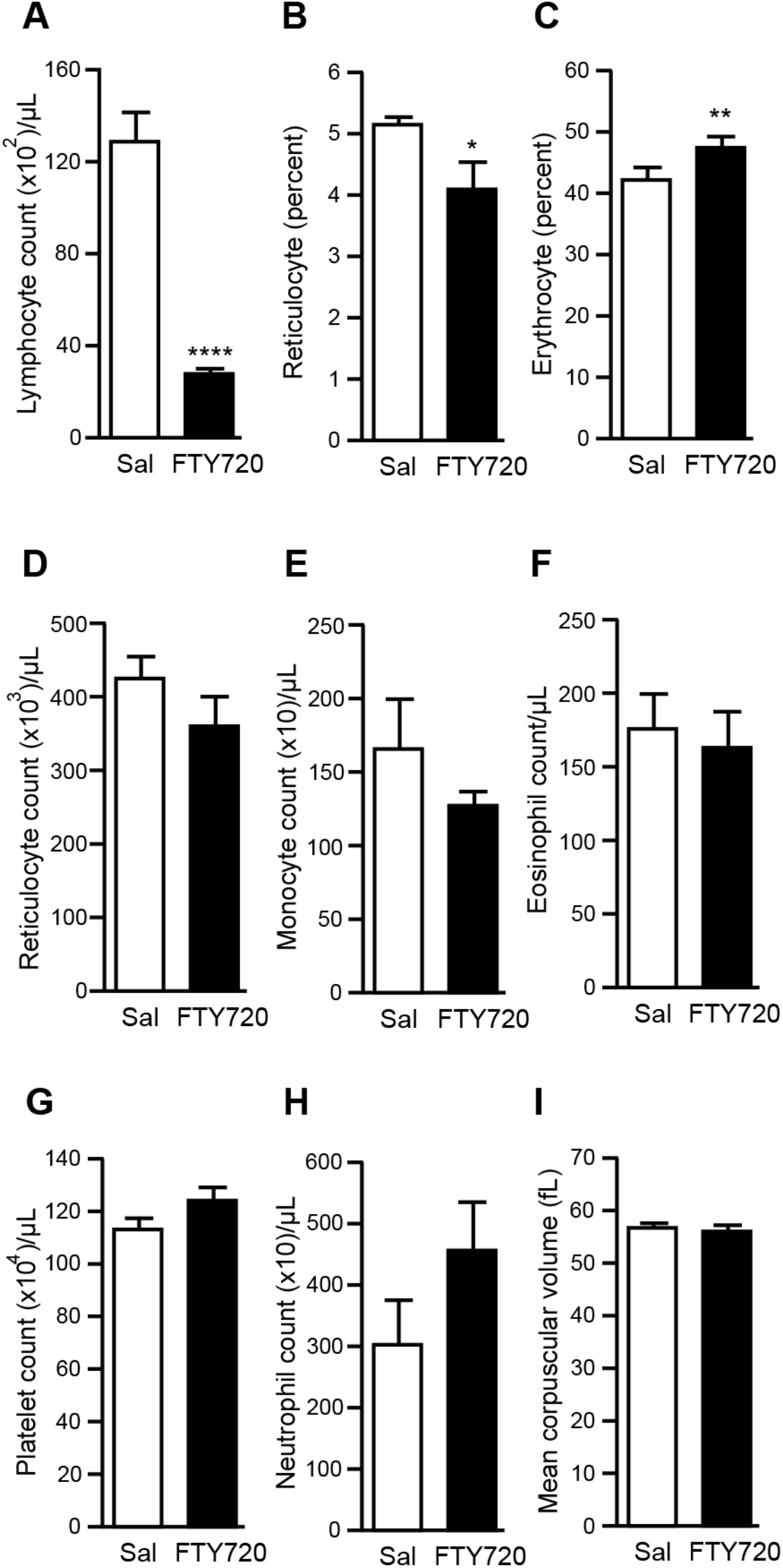
Repeated systemic administration of FTY720 reduced total lymphocyte counts. A) FTY720 reduced total lymphocyte cell counts. B) FTY720 reduced the percentage of immature reticulocytes compared to total reticulocytes. C) FTY720 increased the percentage of the total blood volume composed of erythrocytes. FTY720 did not affect D) reticulocyte count, E) monocyte count, F) eosinophil count, G) platelet count, H) neutrophil count, or I) mean corpuscular volume. Bars represent group means ± SEM; ****p<0.0001, **p<0.01, *p<0.05.

### 3.4. Effects of repeated administration of FTY720 on behaviors

#### 3.4.1. Effects on repeated peripheral FTY720 treatment on behavior in the social interaction test

FTY720 increased the amount of time spent interacting with a stimulus rat in the social interaction test compared to saline-treated rats (t_9_ = 3.884, p < 0.004) (Fig. 3A). This result suggests that FTY720 reduced social anxiety-like behavior.

**Figure 3.**
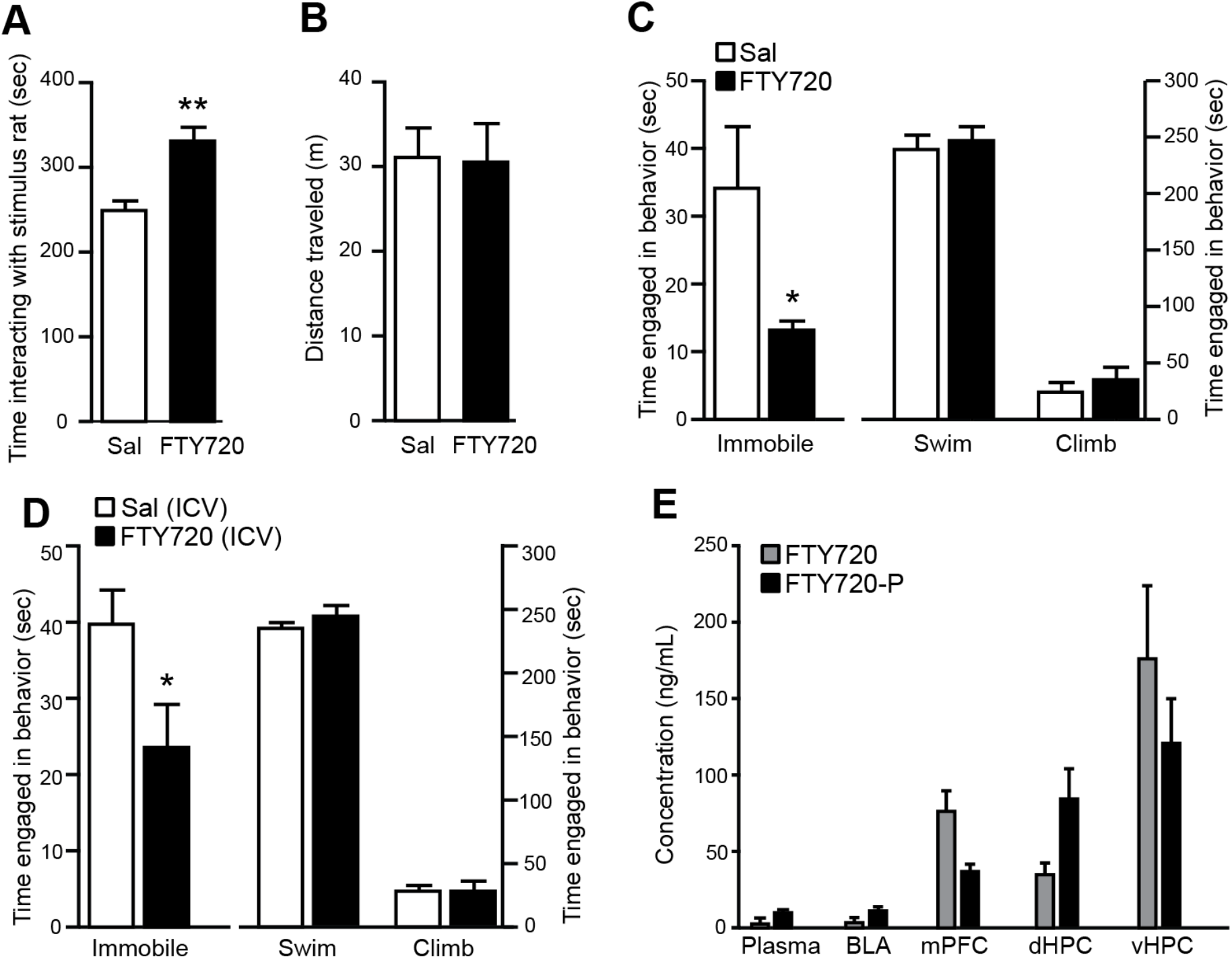
FTY720 reduces behavioral despair and social anxiety-like behavior. A) Intraperitoneal administration of FTY720 increased time interacting with the stimulus rat in the social interaction paradigm. B) FTY720 ip administration did not affect distance traveled in the open field. C) Rats receiving ip injections of FTY720 displayed reduced time spent immobile in the Porsolt forced swim test. Neither time spent swimming or climbing was affected. D) ICV FTY720 treatment reduced time spent immobile in the Porsolt forced swim test. Neither time spent swimming or climbing was affected. E) Concentrations of FTY720 and FTY720-P in plasma and brain regions following ICV delivery of FTY720. Bars represent group means ± SEM; **p<0.01, *p<0.05. BLA, basolateral amygdala; mPFC, medial prefrontal cortex; dHPC, dorsal hippocampus; vHPC ventral hippocampus.

#### 3.4.2. Effects of repeated peripheral FTY720 treatment on behaviors in the open field

No changes in distance traveled were observed as saline- and FTY720-treated rats traveled a similar distance in the open field (t_16_ = 0.43, p < 0.921) (Fig. 3B). This result suggests that FTY720 does not produce any significant effects on locomotor activity

#### 3.4.3. Effects of repeated peripheral FTY720 treatment on behaviors in the Forced Swim Test

FTY720 reduced time spent immobile in the Porsolt FST (t_14_ = 2.42, p < 0.029) compared to saline-treated rats. This result indicates that FTY720 reduced behavioral despair. Time engaged in swimming or climbing were unaffected (Fig. 3C).

#### 3.4.4. Effects of centrally administered FTY720 on behaviors in the Porsolt FST and brain concentrations

We observed reduced time spent in immobility in FTY720-treated rats (t_14_ = 2.27, p < 0.039) (Fig. 3D) with no significant changes in the times spent climbing or swimming, similar to the findings with the ip injection. Following administration of FTY720 into the lateral ventricles, we found that FTY720 and FTY720-P levels were enriched in the mPFC, dorsal (d) and ventral (v) hippocampus (HPC). FTY720 and FTY720-P levels were detected at lower concentrations in the basolateral amygdala (BLA). These data suggest that FTY720 and FTY720-P accumulate in multiple limbic structures following central administration of FTY720. FTY720 and FTY720-P levels were low in blood plasma samples (Fig. 3E), indicating minimal leakage of FTY720 from the brain to periphery.

### 3.5. Effects of repeated peripheral administration of FTY720 on expression of angiogenesis markers in the mPFC

We assessed the expression of 84 angiogenesis- and inflammation-related mRNA transcripts using a targeted PCR array (complete results provided in Table 1). The expression of angiogenesis markers *angiopoietin* (t_13_ = 3.86, p = 0.002), *endothelin 1* (t_13_ = 3.16, p = 0.007), *plasminogen 1* (t_13_ = 2.9, p = 0.012), *vascular endothelial growth factor B* (*Vegf-B*) (t_13_ = 2.21, p = 0.04), and *matrix metalloprotease 2* (*Mmp2*) (t_13_ = 3.09, p = 0.008) were reduced in the mPFC of rats 60min following a third daily injection of FTY720 (Fig. 4A-E). Together, these results provide evidence that FTY720 reduces expression of genes in the mPFC that are involved in angiogenesis. *S1pr1* mRNA was reduced in the mPFC of FTY720-treated rats (t_13_ = 2.25, p = 0.04) (Fig. 4F), indicating that FTY720 reduces transcription of S1PR1.

**Table 1.**
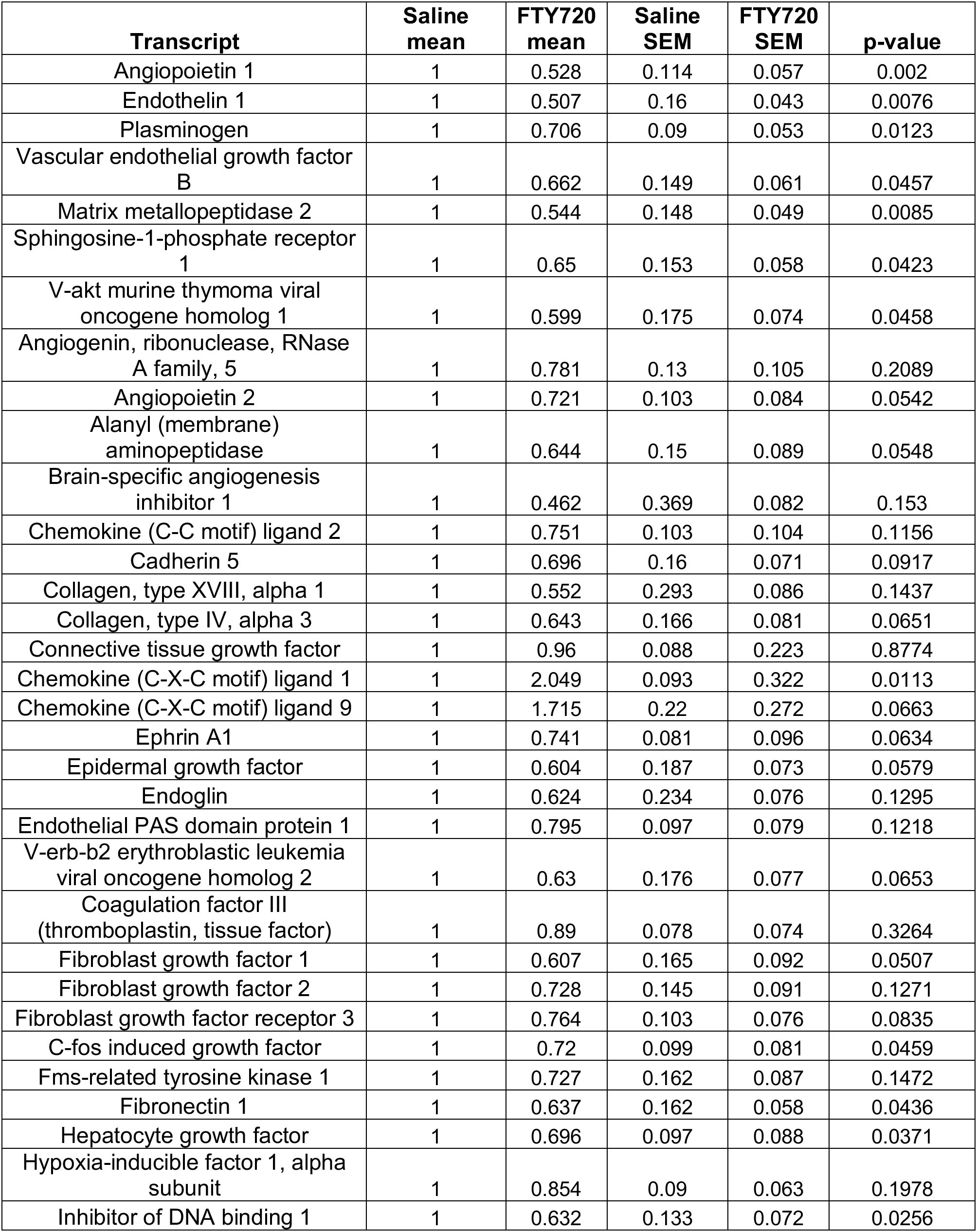
Expression of angiogenesis marker mRNA in the mPFC of saline- and FTY720-treated rats. Data are presented as fold change normalized to controls.

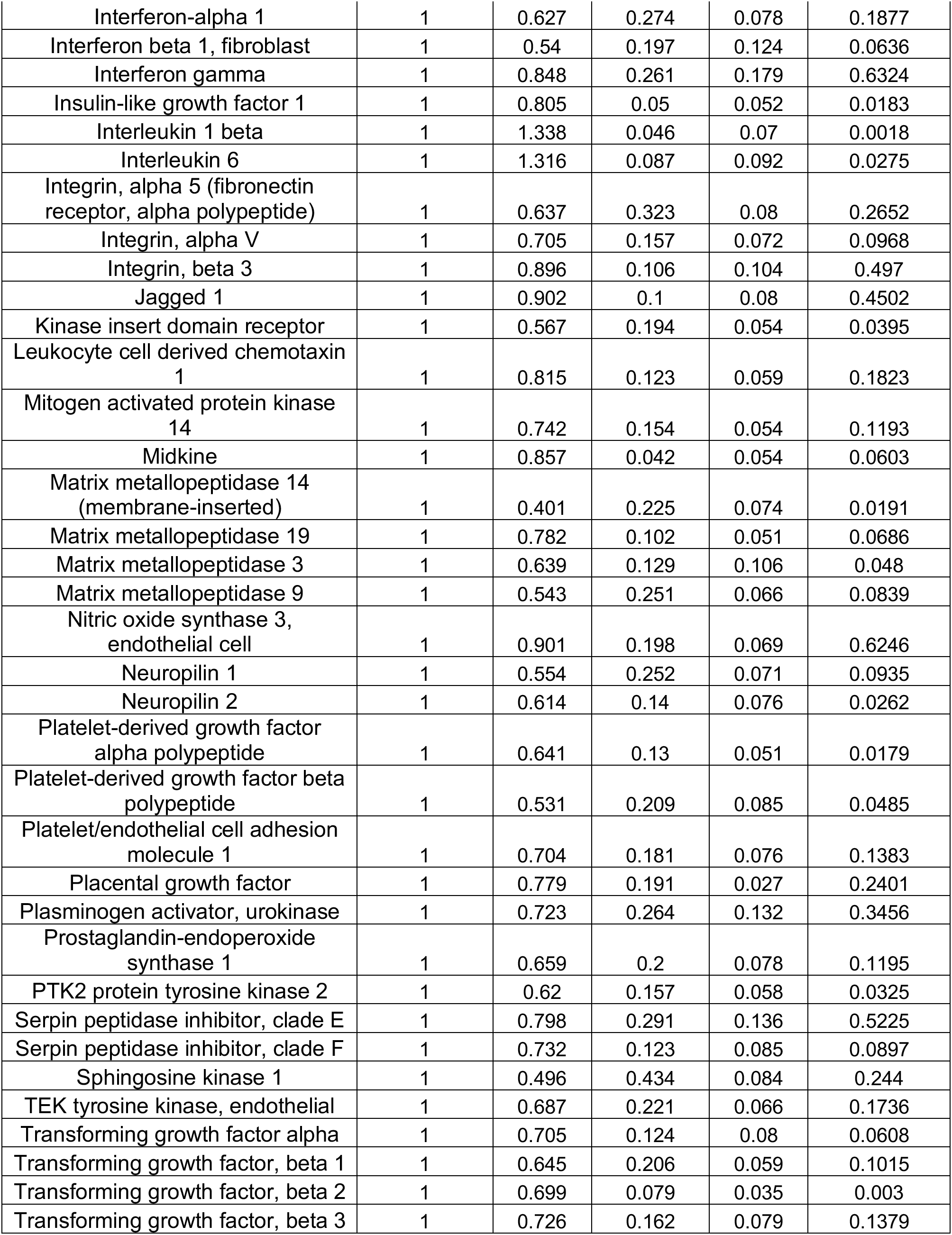

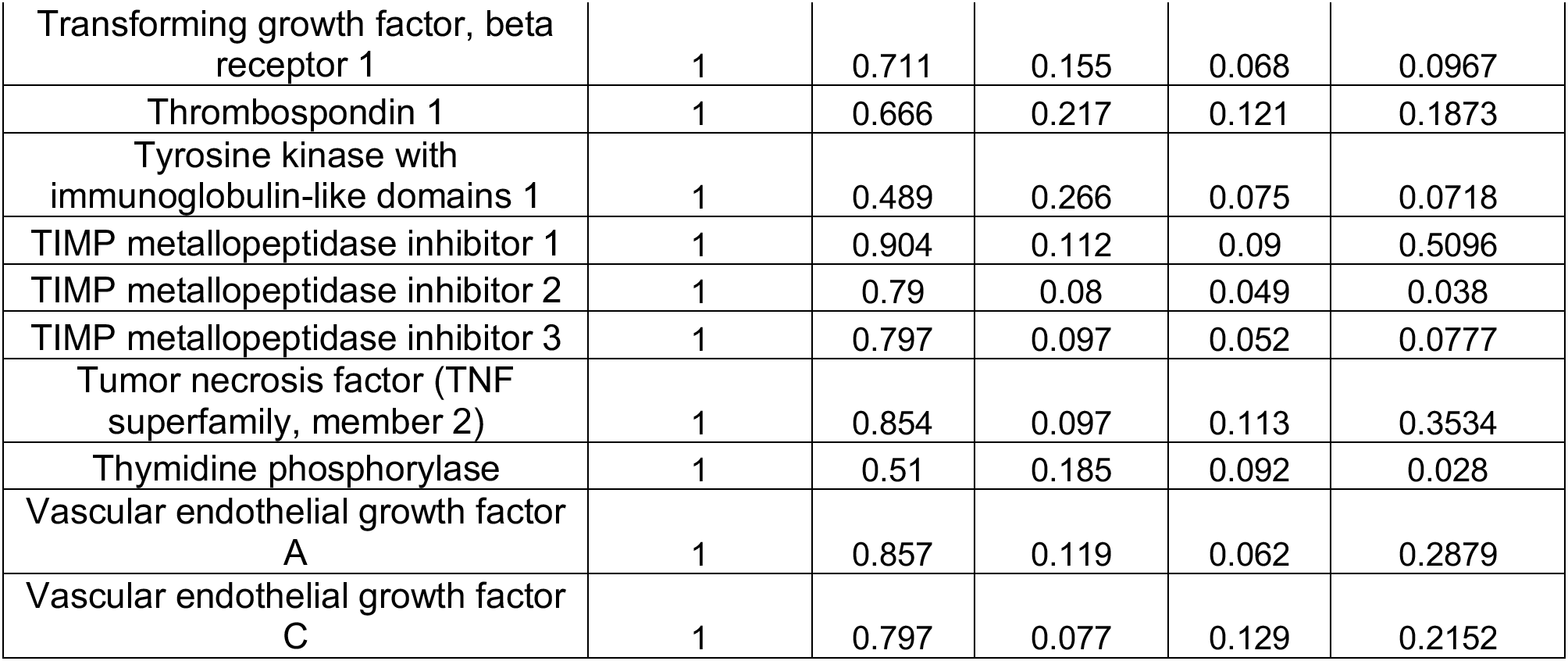

**Figure 4.**
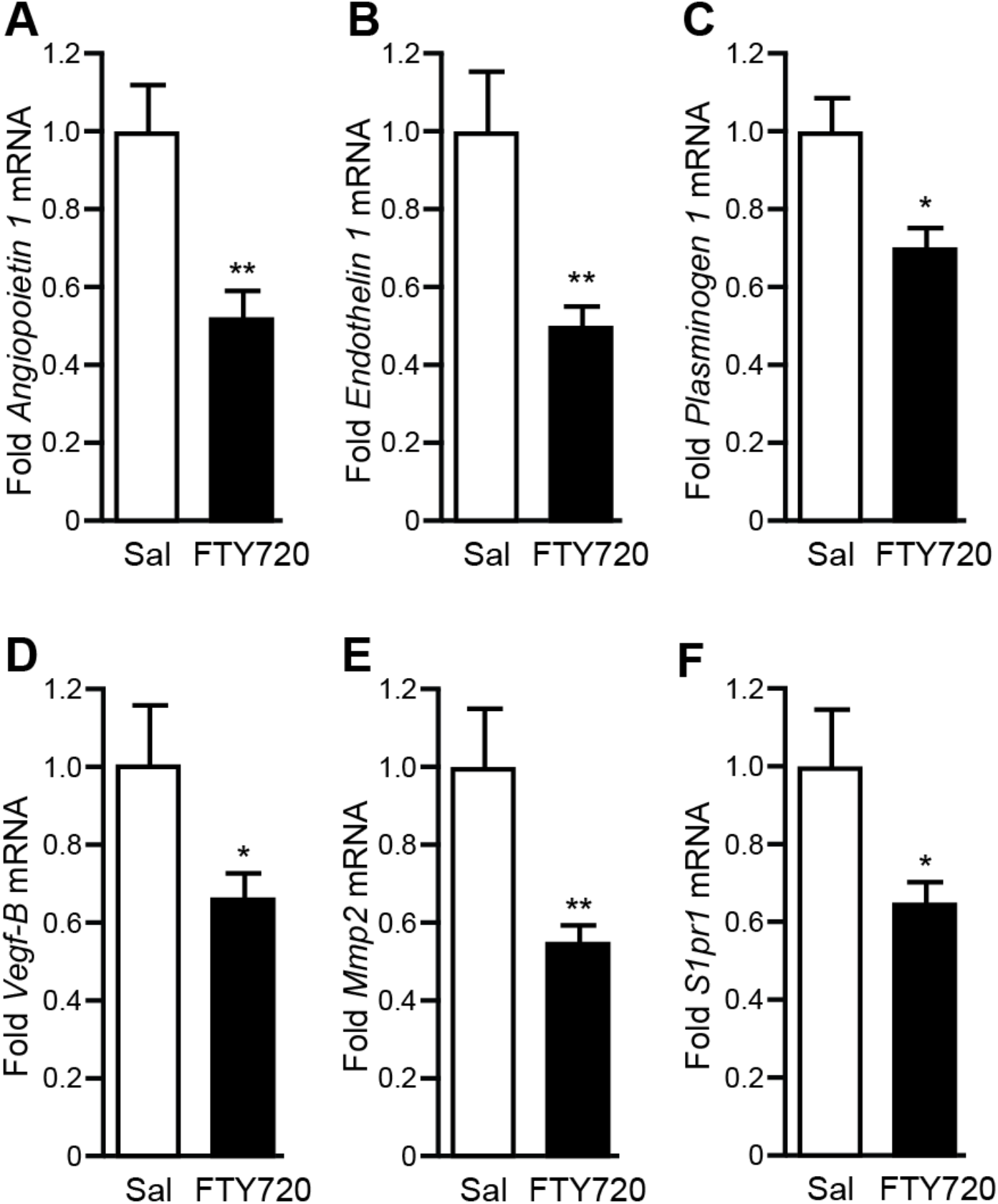
FTY720 treatment alters the expression of genes in the mPFC that regulate angiogenesis. Compared to saline-treated rats, FTY720-treated rats displayed reduced mRNA expression of A) *Angiopoietin 1*, B) *Endothelin 1*, C) *Plasminogen 1*, D) *Vegf-B*, and E) *Mmp2*. F) *S1pr1* mRNA. Bars represent group means ± SEM; **p<0.01, *p<0.05.

## 4. Discussion

In the current study, we determined that FTY720 concentrations increase within 15min of intraperitoneal administration and remain elevated for at least 24 hours. The concentrations of phosphorylated FTY720, FTY720-P, thought to be the physiologically active compound increased more slowly, were elevated at 60min and peaked at 24h after administration. Because of this profile, we examined actions of FT720 administration 60min after a single administration and repeated administration when FTY720 concentrations were expected to be in an elevated steady-state. We observed that FTY720 increased baseline ACTH and corticosterone concentrations. FTY720 administration also reduced lymphocyte counts (ie. lymphopenia) which is typically attributed to impairments in lymphocyte trafficking due to S1PR1 internalization in lymphocytes ^16,17,28^. Thus, because lymphocytes were reduced 60min following a third daily injection we hypothesize that S1PR1 was internalized at this timepoint. However, FTY720 does not internalize all S1PRs ^16^ and therefore we hypothesize that FTY720 was an agonist to other S1PRs at this timepoint. It should be noted that in addition to S1PR1 internalization, elevated glucocorticoids can also induce lymphopenia ^37–40^, and therefore FTY720-induced increases in basal corticosterone concentrations may also contribute to FTY720-mediated lymphopenia. We observed that repeated administration of FTY720 reduced social anxiety-like behavior as assessed by the social interaction paradigm and confirmed previous work that demonstrated antidepressant-like effects of FTY720, which we also observed after icv administration of FTY720. We determined that FTY720 reduced expression of the angiogenesis markers *Angiopoietin*, *Endothelin 1*, *Plasminogen 1*, *Vegf-B,* and *Mmp2* in the mPFC, suggesting that these markers may be important for the behavioral effects of FTY720. We also observed reduced *S1pr1* mRNA expression in the mPFC following FTY720 treatment, suggesting that in addition to known effects of S1PR1 internalization ^16^, FTY720 may further functionally antagonize S1PR1 by reducing their transcription. Together, these results indicate that FTY720 treatment increases baseline HPA axis activity while reducing behavioral despair- and social anxiety-like behavior. Additionally, FTY720 increases angiogenesis markers and reduces *S1PR1* mRNA expression in the mPFC. These results further support a role of S1PRs in the regulation of HPA activity and behaviors related to anxiety and affect.

FTY720 increased baseline ACTH and corticosterone concentrations and produced a blunted HPA response to restraint. Increased baseline corticosterone may contribute to blunted ACTH and corticosterone responses to restraint via fast negative feedback of the stress response ^41,42^. S1PRs are expressed throughout the HPA axis, including the hypothalamus ^43^, the pituitary ^25^, and the adrenal gland ^26^ so it is unclear whether FTY720 induction of basal ACTH and corticosterone is centrally or peripherally mediated. FTY720 could act centrally to impact basal HPA activity. The mPFC is an important regulator of both basal HPA activity ^44,45^ and negative feedback ^46^. FTY720 may act there to regulate basal HPA activity. However, this potential action would have to be mediated through S1PR1 and not through S1PR3 as S1PR3 over-expression or knock-down did not impact basal HPA activity ^8^. S1PRs are thought to be widely distributed in the brain and therefore other sites of actions cannot be ruled out. Alternatively, FTY720 induction of basal ACTH and corticosterone could be mediated by its direct actions in the pituitary and/or adrenal glands. The functions of S1PRs in the pituitary and adrenal glands are not known but would represent a novel mechanism for the regulation of ACTH and glucocorticoids. Basal glucocorticoid concentrations are important for permitting homeostatic metabolic and immune functions and important for optimal functioning of other stress responsive systems ^47^. Basal glucocorticoids are also disrupted in some stress-related psychiatric disorders such as depression ^48^ and PTSD ^49^. Thus, S1PR regulation of glucocorticoid release represents a novel mechanism by which circulating concentrations of these hormones can be titered, ultimately modulating their widespread effects of glucocorticoids on physiology and neural functions under both baseline and stressed conditions. S1PRs could therefore modify a wide variety of metabolic, immune and neural functions through regulation of ACTH or glucocorticoids.

It is possible that shorter durations of FTY720 treatment may increase baseline ACTH and/or glucocorticoids in humans, similar to what we observed here, but there is a report of a lack of effect following long-term treatment of FTY720 on basal cortisol ^50^. One potential mechanism for this lack of effect could be FTY720 inhibition of S1P lyase, the enzyme that degrades S1P ^51^. Loss of function mutations in the gene encoding S1P lyase are associated with cortisol deficiency in certain families ^52,53^ and may be due to adrenal calcification ^54^. The increased baseline plasma concentrations of ACTH and corticosterone reported in this study were observed following relatively short exposure to FTY720 (3 days or less). It is possible that longer exposure to FTY720 could result in adrenal calcification, which may ultimately negate the acute effects of FTY720 in increasing baseline ACTH and corticosterone or due to other mechanisms that have not yet been identified.

Previous work has demonstrated that FTY720 exerts antidepressant-like and anxiolytic effects in rodents ^19^ and humans undergoing treatment for MS ^55^. Here, we confirmed previous preclinical work demonstrating that intraperitoneal injection of FTY720 reduces time spent immobile in the FST and increases time interacting with a stimulus rat in the social interaction paradigm ^19^. We elaborated on these findings by demonstrating that icv injections of FTY720 are sufficient to reduce time spent immobile in the FST. FTY720 administered via icv injection accumulated in the mPFC and hippocampus. Lower levels of FTY720 were observed in the BLA, which may be because of its relatively distal location from ventricles. FTY720 levels were also low in plasma, suggesting that these behavioral effects were due to brain actions of FTY720. For testing in the FST, FTY720 was administered immediately following the training phase and 60min prior to the test phase of the FST. This mimicked the administration regimen that identifies efficacious anti-depressant drugs such as fluoxetine ^36^. FTY720 increases hippocampal BDNF ^20,21^, which provides antidepressant-like effects within a similar timeframe ^22^. The reduced time spent immobile in the FST displayed by FTY720-treated rats may be linked to increased hippocampal BDNF. Thus, FTY720 seems to have anti-depressant properties but further studies are required to determine the mechanisms through which FTY720 acts to reduce behavioral despair and passive behaviors in the FST.

We demonstrated that FTY720 reduces markers for angiogenesis and related processes in the mPFC. Blood vessel plasticity is intimately tied to both neuronal and immune activity. Blood vessels can undergo plasticity and growth to support increases in neuronal activity ^56–59^. We have previously shown that rats vulnerable to repeated social defeat exhibit increases in chronic neuronal activity accompanied by remodeling of blood vessels in the ventral hippocampus ^32^. Blood vessels can also become permeable resulting in leakage of peripheral immune or other factors that increase microglial activity which promotes expression of pro-inflammatory cytokines ^60,61^. In the mPFC, we previously showed that S1PR3 signaling (FTY720 is a partial agonist for S1PR3 ^16^) reduces pro-inflammatory cytokine expression and promote resilience to social defeat, by reducing immobility in the FST and social anxiety ^8^. Together, these findings suggest that FTY720 actions through processes tied to blood vessel remodeling and angiogenesis may underlie the reductions in behavioral despair and social anxiety we observed with FTY720 administration.

We identified decreases of *angiopoietin 1*, *endothelin 1*, *plasminogen 1*, *Vegf-B*, and *Mmp2* in the mPFC 60min following a third daily injection of FTY-720. Angiopoietin 1 binds to tyrosine protein kinase receptors on endothelial cells and mediates angiogenesis through reciprocal interactions between endothelial cells and the extracellular matrix ^62^. Endothelin 1 binds to the endothelin_A_ G-protein coupled receptor to induce angiogenesis and vasoconstriction ^63^. Plasminogen 1 is cleaved to become active plasmin, a protease involved in the cleavage of blood plasma proteins including fibrin, which mediates blood clotting ^64^. Vegf-B binds to Vegf receptor 1 and is critical for blood vessel survival. Vegf-B is not critical for blood vessel growth under normal conditions but is critical for blood vessel growth under pathological conditions ^32,65^. Mmp2 regulates angiogenesis by promoting breakdown of the extracellular matrix, a process critical for tissue remodeling and angiogenesis ^66^. Thus, FTY720 reduces the expression of multiple genes important for angiogenesis. This may decrease angiogenesis through impaired blood vessel survival and reduced remodeling of the extracellular matrix.

Glucocorticoids are generally reported to induce depressive- and anxiety-like behavior ^67^. Thus, the effects of FTY720 on increasing ACTH and corticosterone may seem contradictory to its effects on reducing despair- and social anxiety-like behavior. One possible explanation is that FTY720-mediated reductions in despair- and social anxiety-like behaviors are attributed to the central mechanisms described above and function independently of peripheral effects on HPA axis activity. Therefore, while FTY720-induced increases in corticosterone may promote despair- and anxiety-like behavior on their own, these effects may be overcome by central effects of FTY720 that reduce despair- and social anxiety-like behavior. Further studies are required to determine the specific S1PRs and the target sites that mediate these effects of FTY720.

Regardless of the site of action, our results contribute to our growing understanding of S1PR functions and the results here suggest that S1PRs are novel regulators of baseline HPA axis activity and behaviors relevant to affect and anxiety.

This work was supported by the Defense Advanced Research Projects Agency (DARPA) and the U.S. Army Research Office under grant number W911NF1010093 to SB. Additional support in the form of a Training Grant in Neurodevelopmental Disabilities NIH/NINDS T32 NS007413 and a NARSAD Young Investigator Grant from the Brain & Behavioral Research Foundation (29185) was awarded to BC. The authors have no conflicts of interest.

